# Individual differences show that only some bats can cope with noise-induced masking and distraction

**DOI:** 10.1101/2020.07.04.188086

**Authors:** Dylan G.E. Gomes, Holger R. Goerlitz

**Affiliations:** Acoustic and Functional Ecology, Max Plank Institute for Ornithology, Seewiesen, Germany; Boise State University, Boise, Idaho, USA, 83725-1515

**Keywords:** anthropogenic noise, echolocation, mechanism, *Chiroptera*, *Phyllostomidae*, discrimination task

## Abstract

Anthropogenic noise is a widespread pollutant that has received considerable recent attention. While alarming effects on wildlife have been documented, we have limited understanding of the perceptual mechanisms of noise disturbance, which are required to understand potential mitigation measures. Likewise, individual differences in response to noise (especially via perceptual mechanisms) are likely widespread, but lacking in empirical data. Here we use echolocating bats, a trained discrimination task, and experimental noise playback to explicitly test perceptual mechanisms of noise disturbance. We demonstrate high individual variability in response to noise treatments and evidence for multiple perceptual mechanisms. Additionally, we highlight that only some individuals are able to cope with noise, while others are not. We tested for changes in echolocation call duration, amplitude, and peak frequency as possible ways of coping with noise. Although all bats strongly increased call amplitude and showed additional minor changes in call duration and frequency, these changes cannot explain the differences in coping and non-coping individuals. Our understanding of noise disturbance needs to become more mechanistic and individualistic as research knowledge is transformed into policy changes and conservation action.

## Introduction

Anthropogenic noise is a global pollutant that has recently gained considerable attention by behavioral biologists (Barber et al., 2010). Noise can disrupt animal behavior, such as communication (Brumm and Slabbekoorn, 2005; Rabin et al., 2003) and foraging (Gomes et al., 2016; Purser and Radford, 2011; Siemers and Schaub, 2011), reduce reproductive success (Halfwerk et al., 2011), increase mortality (Simpson et al., 2016), change biological communities (Francis et al., 2011), and alter ecological services (Francis et al., 2012). Yet it is not often understood what mechanisms drive these changes, and if and how different individuals are affected by these mechanisms differently. Individual differences in response to noise has been documented in humans (Furnham and Strbac, 2002; Standing et al., 1990), birds (Naguib et al., 2013), fish (Bruintjes and Radford, 2013) and mongooses (Eastcott et al., 2020), among many others (reviewed in Harding et al., 2019), yet this is often overlooked as individuals are grouped together for analysis. Similarly, researchers rarely test mechanisms of noise disturbance (but see (Zhou et al., 2019). Understanding how we may be able to mitigate the consequences of noise relies heavily on knowledge of direct mechanisms of noise disturbance on individuals. Dominoni et al. (2020), for example, highlight three main perceptual mechanisms of noise disturbance – masking, distraction, and misleading. While these mechanisms apply to all senses, we here consider them specifically in the auditory domain.

Masking is a mechanism whereby noise overlaps in frequency with important signals or cues, thus making the detection and auditory analysis of the signal difficult, if not impossible (Clark et al., 2009; Fay and Wilber, 1989; Gomes et al., 2016; Tanner Jr, 1958). Distraction, on the other hand, occurs when noise competes for the finite attention of an organism, and is not limited to frequencies that overlap with a signal or cue of interest (Chan et al., 2010). Misleading occurs when noise is interpreted as something that it is not, also termed a false alarm (Wiley, 2013), for example a predator (Tyack et al., 2011) or something unknown that might be dangerous. Other mechanisms of disturbance have been proposed, such as stress, fear, and avoidance (Campo et al., 2005; Luo et al., 2015a; Voellmy et al., 2014), yet these physiological and behavioral responses must occur downstream of the initial perceptual mechanism (i.e. masking, distraction, or misleading).

Here, we use a behavioral experiment to tease apart the relative contribution of both masking and distraction as perceptual mechanisms on individual echolocating bats. Echolocating bats are a worthwhile system to study these questions because they actively sense their world via sound. Thus, we can directly interfere with their ability to perceive objects in their environment, and we can track the animals’ efforts and sensory strategies to change and improve their perception relatively easily. That is, we can monitor changes to echolocation call characteristics as a way to understand how these animals are responding to various stimuli.

We trained bats to discriminate surface structures with increasing level of difficulty and under three noise treatments. We made distinct predictions for each of the tested perceptual mechanisms. First, by broadcasting noise that does and does not spectrally overlap with echolocation calls, we directly tested the role of masking. We predicted that masking should only reduce the discrimination performance for spectrally overlapping noise, but not for non-overlapping noise. Second, by testing discrimination performance for tasks of increasing difficulty, we tested the role of distraction. Since distraction assumes that deficits result from limited attentional resources, we predicted that noise-mediated distraction should lower discrimination performance as tasks become more difficult. We predicted further that distraction should be independent of the noises’ spectral overlap with echolocation calls (distinguishing it from masking), but should depend on the noises’ temporal structure. We thus also presented a spectrally overlapping ‘sparse’ noise with random temporal gaps, making the noise less predictable, and thus, more distracting (Glass and Singer, 1972; Kjellberg et al., 1996; Matthews et al., 1980). At the same time, sparse noise might allow bats to listen in-between the noise gaps (“dip listening”), reducing its masking effect (Vélez and Bee, 2011). Thus, if distraction is the primary mechanism of disturbance, then sparse noise should decrease discrimination performance and increase trial duration, while the opposite should be true if masking is the primary mechanism of disturbance.

## Materials and methods

### Animal Husbandry

A captive colony of lesser spear-nosed bats (*Phyllostomus discolor*; Wagner, 1843) were kept in a temperature (~25 °C) and humidity (~70%) controlled room at the Max Planck Institute for Ornithology, Seewiesen, Germany, where they had access to water *ad libitum*, and were fed a fruit-based diet. During experimental days, bats were first only fed during experiments (mealworm reward; *see below*), to maintain motivation. At the end of the day, several hours later, bats were fed fruit. Experiments were carried out in a nearby, but separate room (~21 °C / 65% hum). Bat housing and all research was approved by the German authorities under the permit numbers 311.5-5682.1/1-2014-023 (Landratsamt Starnberg) and 55.2-1-54-2532-18-15 (Regierung von Oberbayern), respectively.

### Experimental Setup

Experiments were conducted in a dark chamber within a dark room (see below for light levels). Walls of both the chamber and the room were covered in anechoic foam to reduce echoes. The chamber held a custom-built mushroom maze (87 cm x 65 cm x 18 cm, W x H x D; (Baier et al., 2019), which allowed the bats perceptual access to two simultaneously presented stimulus discs (reference plus test disc) on either side of the maze (**Figure 1A**). One infrared light barrier next to each of the disc positions objectively recorded the choice of the bat via a custom-written Matlab code (The Mathworks, Nattick, MA, USA), avoiding observer bias and potential observer errors. Two speakers (Vifa, Peerless by Tymphany, San Rafel, CA, USA; power amplifier: TA-FE330R, Sony, Tokyo, Japan) were mounted on either side of the setup for noise playback (**Figure 1A**). The experimenter (stationed outside of the chamber) observed the experiment via a red-filtered computer screen displaying a live-feed from an infrared camera (Foculus FO432SB; NET-GmbH, Finning, Germany; 880 nm infrared LED-illumination, TV6818; ABUS, Wetter, Germany).

**Figure 1:**
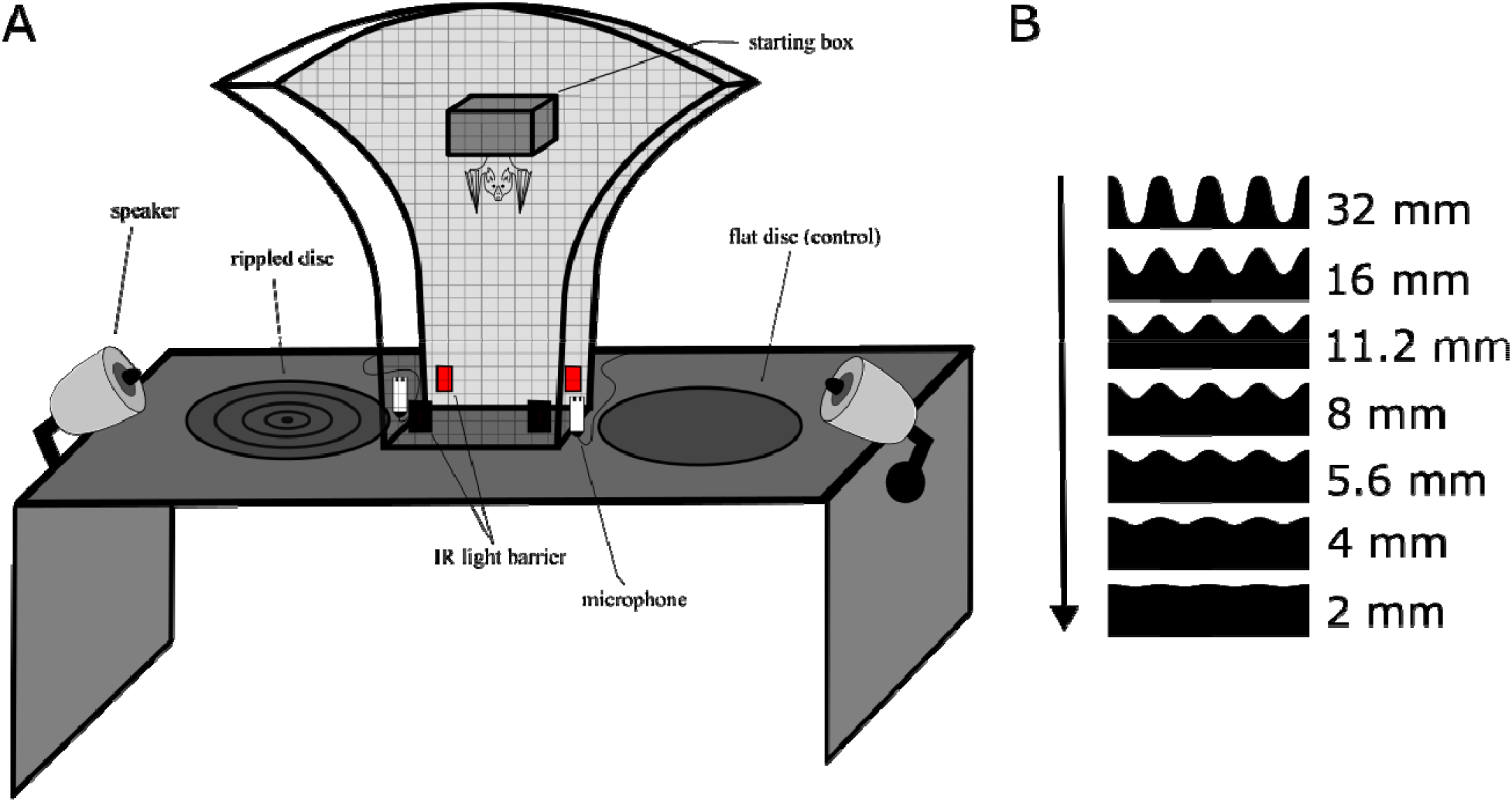
Sketch of the experimental setup. A) The mushroom-shape of the maze allowed the bat simultaneous perceptual access to both discs from multiple angles. Each trial started when the bat left from within the starting box and ended when the bat crossed an infrared light barrier next to each disc, objectively determining decision and duration of each trial. The bat received a food reward for approaching the flat reference disk. Noise treatments were presented via two speakers from similar directions as the returning disc echoes, echolocation call were recorded via microphones next to the light barriers. B) Stimuli used in the discrimination experiment. Cross-section of the stimulus discs and are scaled to size; peak-to-peak ripple height is indicated. As ripple height gets smaller, the task to discriminate the rippled disk from the flat reference disc becomes more difficult, as indicated by the arrow.

### Stimuli

We used an established behavioral assay that has been previously used to test perceptual performance in bats (Baier et al., 2019). We used eight discs with 45 cm diameter as physical stimuli. The stimulus discs were made by a milling cutter (Modellbau Grossmann, Calw, Germany) and then spray-painted with multiple coats to be smooth-textured. One disc (“reference disc”), had a completely flat surface. The seven other discs had concentric ripples, resembling concentric sinusoidal standing waves. All rippled disc had the same spatial frequency of 17.8 ripples per meter, corresponding to eight full sinusoidal ripples per disc, but different ripple heights increasing from 2 to 32 mm peak-to-peak height (2, 4, 5.6, 8, 11.2, 16, 32 mm; **Figure 1B**).

### Noise Treatments

In addition to silence, used as a control, we tested the bats under three white noise treatments: 1) Smooth non-overlapping noise: band-limited white Gaussian noise not overlapping in frequency with the echolocation calls of P. discolor, ranging from 5-35 kHz (10^th^-order butterworth filter). 2) Smooth-overlapping noise: band-limited white Gaussian noise overlapping in frequency with the echolocation calls of *P. discolor*, ranging from 40-90 kHz (10^th^-order buttherworth filter). 3) Sparse-overlapping noise: a temporally fluctuating overlapping noise based on the smooth-overlapping noise (40-90 kHz), where silent gaps with a uniformly random duration (mean: 0.3 ms, range: 0-0.6 ms) were inserted between adjacent samples (Hübner and Wiegrebe, 2003). This generates temporal fluctuations in the temporal envelope of the noise, causing the noise to sound rougher in comparison to the smooth noise. This is quantified by the base-10 logarithm of the fourth moment (Hartmann and Pumplin, 1988), which was 1.44 logM4 compared to 0.48 logM4 for the two smooth noises (cf. Grunwald et al., 2004). We initially generated uncorrelated stereo noise files of 60 min duration and corrected each channel for the corresponding speaker’s frequency response (Matlab). Noise playback was broadcast continuously at 70 dB SPL RMS re. 20 μPa at the starting position of the bat throughout each bat’s daily experimental session, starting at least 30 s prior to the beginning of the first trial. It is important to note that the perceived loudness was likely different, since the noise treatments had different bandwidths and the auditory sensitivity of *P. discolor* varies over their range of hearing (Esser and Daucher, 1996; Hoffmann et al., 2008). However, this should not affect any interpretation of the designed tests of masking and distraction.

### Training and testing

Four bats (*P. discolor*) were trained in a two-alternative forced-choice paradigm to discriminate the flat reference disc from the stimulus disc with the highest ripples (32 mm). During training, bats received mealworms (larvae of *Tenebrio molitor*) as reward when approaching the flat reference disc only. Once bats consistently approached the flat reference disc (>70% of the trials) during three consecutive days, they were considered trained and data acquisition started. Throughout testing and training, the flat reference disc was pseudo-randomly (Gellermann, 1933) alternated between each side of the experimental setup to avoid any biases in location preferences by the bats.

Prior to each trial, bats were encouraged to enter a small Tupperware ‘starting’ box in the middle of the experimental setup by offering a blended banana food reward via a syringe tube that was mounted inside this starting box (**Figure 1A**). While in the starting box, bats had no perceptual access to the discs since the solid bottom door was closed upon entering and the discs were swiveled to their positions. Once the starting box opened, the trial started. Bats were allowed to crawl through the setup towards the discs. When they broke the IR light barrier, the trial ended, and the bat was rewarded with a mealworm when it chose the flat reference disc.

Bats were initially tested in a silent (*i.e.* ambient sound level) experimental room to generate baseline psychometric curves with discs of 5 different ripple heights (2 mm, 4 mm, 8 mm, 16 mm, and 32 mm). For the subsequent tests, we added two additional discs with intermediate ripple heights (5.6 mm and 11.2 mm) to get better resolution around the turning point of the psychometric function measured in silence. The bats were then tested with all seven discs in each of the three noise conditions in pseudo-random sequence. Finally, each bat was retested a final time in silent conditions to assure that differences in performance were not due to learning or other order effects (all discs). Each bat was tested 30 times for every ripple height and noise treatment combination, totaling 990 trials per individual bat.

To motivate the bats, each day was started with easier discrimination tasks (higher ripple heights) and gradually moved towards more difficult tasks (lower ripple heights). Bats were allowed to continue testing until satiated or no longer food-motivated, which was determined by the bat attempting to leave the mushroom maze via an exit door in the top of the setup.

### Aborted trials

If the bat did not exit the starting box within 5 minutes after starting a trial, the trial was aborted and repeated. As the bats did not make a decision in those aborted trials, they were not included in further analyses. The one exception, however, is that we analyzed the number of these aborted trials as a measure of aversion to the noise. This was behaviorally distinguished from satiation, as bats would crawl toward the door to exit the maze when they were seemingly no longer food-motivated.

### Echolocation call recording and analysis

We recorded the bats’ echolocation calls during the four seconds prior to the decision of each trial, using two microphones (Knowles SPU0410) positioned just behind each light barrier, a sound card (Fireface 802, RME, Haimhausen, Germany; 192 kHz sampling rate, 16-bit resolution) and playrec (V2.1.0, playrec.co.uk) for Matlab (V2007b, The Mathworks, Nattick, MA, USA).

Echolocation calls were analyzed automatically by custom-written scripts in Matlab (V2016a, The Mathworks, Nattick, MA, USA), advanced from previous work (Goerlitz et al., 2008; Luo et al., 2015a). First, we filtered all recordings with each microphone’s compensatory impulse response (511-order finite impulse response filter) to compensate for the microphone’s frequency response, and a band-pass filter (38-95 kHz, 8th-order elliptic filter). Second, we used a threshold detector to broadly determine the timing of all acoustic events: we additionally band-pass-filtered recordings around the bats’ main call energy (45-90 kHz, 4th-order elliptic filter), calculated their low-pass filtered (500 Hz, 4th-order elliptic filter) Hilbert-envelope, and detected all acoustic events that surpassed a threshold (mean + 2x STD of the envelope), excluding events that were too close to the preceding event (< 20 ms) and too short (< 0.75 ms). We then added an additional 0.5 ms on both sides of the detected acoustic events, which, together with the previous low-pass-filtering of the envelope, ensured that the determined time window included the full call flanked by non-call samples. Third, we detected the actual call within this time window of the recording, and analyzed its acoustic parameters.

Call duration was determined from the low-pass filtered (5000 Hz, 2nd-order butterworth filter) Hilbert envelope of the originally filtered recording (38-95 kHz) at −12 dB below the envelope’s peak value. Peak frequency (frequency with highest amplitude), frequency centroid (dividing the call energy into two halves along the frequency axis; Au, 2012) and the lowest and highest frequency (frequencies with amplitudes −12 dB below the highest amplitude) were calculated from the time-averaged call spectrogram (1024 FFT of 100 samples, 95% overlap). Relative call level was calculated as the root mean square (RMS) of all samples within the −12 dB duration criterion and expressed in dB FS, i.e., negative dB values relative to the full scale of the recording system.

If a call was detected on both microphones, we only analyzed the call with the higher signal-to-noise-ratio (SNR: call-RMS relative to RMS of all parts of the recording that were not classified as acoustic events). Of all recorded calls (N = 287,061), we excluded for further analysis calls shorter than 0.3 ms and longer than 2 ms, with too high (>−0.5 dB FS, to avoid clipping) or too low recorded peak amplitudes (<−15 dB FS), with a SNR of less than 20 dB, and whose ratio between the −12 dB duration and the −6 dB duration was larger than 1.5 (to exclude calls with long echoes). All remaining calls (N = 63,990) were manually viewed as spectrogram (256 FFT, 50 time slices over full call length, 95% overlap), blind to bat individual and noise treatment, to exclude ambiguous recordings and obvious artefacts, e.g., overlapping call-echo-pairs and non-multiharmonic sounds (e.g., clicks, external noise), resulting in a final data set of 59,173 calls (0-83 calls per trial, 3469 trials (of 3960 trials in total, 87.6%) with at least one call, 3166 trials (80.0%) with at least 3 calls, 2912 trials (73.5%) with at least 5 calls, 2265 trials (57.2%) with at least 10 calls. For further analysis, we used the mean (grouped by each trial) call parameters in statistical models. Note that the background noise did not affect our call level measurements because we only analyzed calls with a SNR > 20 dB.

### Visual system and light levels

Light levels in the experimental room were extremely low (1.39 × 10^−5^ lux; SPM068 with ILT1700 light detector, resolution 10^−7^ lux, International Light Technologies, Peabody, MA, USA), precluding the use of vision to discriminate between discs. Many other laboratory experiments, which have similarly excluded the use of vision due to an assumed unavailability of light, have either reported higher light levels than us or did not measure or report light levels. Additionally, it has been experimentally shown that another related Phyllostomid bat (*Macrotus californicus*) only has visual acuity to light levels as low as 2 × 10^−3^ lux (Bell and Fenton, 1986), which is nearly two orders of magnitude higher than our light levels. Furthermore, *M. californicus* has one of the highest sensitivities to low light levels known (Bell and Fenton, 1986; Eklöf et al., 2014). Thus, it is extremely unlikely that the *Phyllostomus discolor* used here were able to visually discriminate between the discs.

### Statistical analysis

We fitted (generalized) linear models to the behavioral data of each individual, using R (R Core Team, 2017). Response variables were analyzed with different distribution families and link functions based on theoretical sampling distributions of the data, and model fits were validated with plots of model residuals, and were checked for collinearity.

We used a binomial distribution family and logit link function to analyze differences in discrimination performance and number of aborted trials, since these were binary data. We used an inverse Gaussian distribution family with an identity link function to analyze trial time data (Baayen and Milin, 2010). Log-normal linear models (Gaussian family with an identity function) were used to analyze log-transformed received call level, duration, peak frequency, and frequency centroid. Peak frequency and frequency centroid measure similar aspects of vocalization frequency and are both used in the literature (Goerlitz et al., 2008; Holderied et al., 2005; Lattenkamp et al., 2018; Lazure and Fenton, 2011). We therefore included both metrics in our analyses for comparability, but report only peak frequency in the main text because it is the most commonly used metric, and present frequency centroid data in the supplementary information (**Table S6**).

For all models we used noise treatment, ripple height, and their interaction as explanatory variables, while the number of days that bats were in our experiment was included as a covariate – all fitted as fixed effects. The number of days an animal is in an experiment may be important because animals can learn over time to be faster at a given task, or, conversely, they become frustrated with difficult tasks. Including this term in our model allowed us to account for this, while being able to make inferences about any patterns that emerge.

We fitted individual models for each bat, instead of single models for every response variable, with bats as random effects terms, for two reasons. Firstly, it has been suggested that random effects terms should have a minimum of five groups; otherwise estimates of variance become imprecise (Harrison et al., 2018). As we only had 4 bats complete the experiment, we were unable to fill this requirement. Secondly, and more importantly, fitting models to each individual bat allowed us to understand the nuanced differences between them, which an all-bats-combined model would not achieve. Since we fitted four models per response variable (one for each bat), we used conservative Bonferroni corrections to correct *p* values for these multiple comparisons by multiplying *p* values by 4. All differences reported in results due to noise treatments are model estimates, and not differences in raw data.

### Performance thresholds

We used a binomial generalized linear model with a probit link (constrained between 0.5 and 1 with the link function `mafc.probit` in the R package `psyphy`; Knoblauch, 2007) to generate estimates of ripple height thresholds at which bats exceed correct responses at least 70% of the time. For each bat, 1000 simulated discrimination performance (0 or 1) datasets were generated based on the above model estimates for each bat, at each ripple height, within each noise treatment. Then the lower 0.025 and upper 0.975 percentiles of those data gave us a 95% confidence interval band around our performance threshold.

## Results

### Discrimination performance

All four bats learned to discriminate the smooth disc from the rippled disc with the highest ripples (32 mm) in silence (88-100% correct), and showed reduced discrimination performance with decreasing ripple height (**Figure 2**, orange line; logistic model *p* < 0.001; **Table 1**). In silence, performance dropped below our 70% threshold criterion for ripple heights around 7.9 mm (mean; range = 5.4 – 11.4 mm; **Figure 2; Table 2**), matching the mean threshold found by (Baier et al., 2019) of 8.0 mm (range: 3.7 – 12.3 mm). Noise treatments did not change the discrimination performance of bats A and B (hereafter ‘coping’ bats; **Table 1**), and the 95% confidence intervals of their thresholds in noise overlapped with those in silence. In contrast, discrimination performance decreased for bats C and D (hereafter ‘non-coping’ bats) both under smooth-overlapping (z = −3.2, *p* < 0.01; z = −3.2, *p* < 0.01) and sparse-overlapping (z = −3.7, *p* < 0.001; z = −3.1, *p* < 0.01) noise (blue and purple lines in **Figure 2** respectively). The same is true for the smooth non-overlapping noise for bat C (z = −2.5, *p* < 0.05), yet not for bat D (z = 1.8, *p* = 0.24).

**Table 1:** Result of a generalized linear model for discrimination performance in various noise treatments. Model results show the estimated differences in discrimination performance (relative to control trials) for the three noise treatments, ripple height, the number of days the bat was in the experiment, and the interaction between each noise treatment and ripple height (i.e. the shape of each performance curve as a function of ripple height), separately for each bat. Data were analyzed with binomial distribution and logit link function.

| Bat | Variable | Estimate | SE | Z value | p value |
| --- | --- | --- | --- | --- | --- |
| A | (Intercept) | -0.766 | 0.275 | -2.791 | 0.02 |
| A | Non-overlapping | 0.228 | 0.521 | 0.438 | 0.987 |
| A | Overlapping | 0.317 | 0.423 | 0.749 | 0.911 |
| A | Sparse OL | 0.544 | 0.447 | 1.216 | 0.637 |
| A | Ripple height | 0.294 | 0.047 | 6.222 | <0.001 |
| A | Day of experiment | 0.110 | 0.095 | 1.165 | 0.673 |
| A | Non-overlapping:Ripple height | 0.079 | 0.099 | 0.798 | 0.891 |
| A | Overlapping:Ripple height | -0.096 | 0.064 | -1.504 | 0.435 |
| A | Sparse OL:Ripple height | -0.088 | 0.069 | -1.265 | 0.603 |
| B | (Intercept) | -0.544 | 0.243 | -2.240 | 0.096 |
| B | Non-overlapping | 0.200 | 0.412 | 0.486 | 0.981 |
| B | Overlapping | 0.441 | 0.377 | 1.171 | 0.668 |
| B | Sparse OL | 0.490 | 0.364 | 1.346 | 0.543 |
| B | Ripple height | 0.216 | 0.036 | 6.082 | <0.001 |
| B | Day of experiment | 0.003 | 0.107 | 0.025 | 1 |
| B | Non-overlapping:Ripple height | -0.012 | 0.059 | -0.209 | 0.999 |
| B | Overlapping:Ripple height | -0.091 | 0.048 | -1.905 | 0.209 |
| B | Sparse OL:Ripple height | -0.109 | 0.045 | -2.405 | 0.062 |
| C | (Intercept) | -0.130 | 0.200 | -0.649 | 0.945 |
| C | Non-overlapping | 0.277 | 0.312 | 0.888 | 0.847 |
| C | Overlapping | 0.228 | 0.305 | 0.745 | 0.912 |
| C | Sparse OL | 0.433 | 0.298 | 1.451 | 0.471 |
| C | Ripple height | 0.118 | 0.022 | 5.465 | <0.001 |
| C | Day of experiment | -0.071 | 0.087 | -0.817 | 0.882 |
| C | Non-overlapping:Ripple height | -0.071 | 0.028 | -2.518 | 0.047 |
| C | Overlapping:Ripple height | -0.085 | 0.027 | -3.157 | 0.008 |
| C | Sparse OL:Ripple height | -0.100 | 0.027 | -3.726 | <0.001 |
| D | (Intercept) | -0.069 | 0.177 | -0.387 | 0.992 |
| D | Non-overlapping | -0.486 | 0.327 | -1.487 | 0.445 |
| D | Overlapping | 0.118 | 0.278 | 0.422 | 0.989 |
| D | Sparse OL | 0.280 | 0.283 | 0.990 | 0.789 |
| D | Ripple height | 0.074 | 0.015 | 4.937 | <0.001 |
| D | Day of experiment | 0.085 | 0.076 | 1.129 | 0.699 |
| D | Non-overlapping:Ripple height | 0.060 | 0.033 | 1.832 | 0.242 |
| D | Overlapping:Ripple height | -0.066 | 0.021 | -3.179 | 0.004 |
| D | Sparse OL:Ripple height | -0.066 | 0.021 | -3.141 | 0.008 |

**Table 2:** Threshold of the discrimination performance for ripple detection. The threshold is the ripple height where bats exceeded a 0.7 probability of a correct choice. For each bat, 1000 simulated discrimination thresholds were generated with a binomial generalized linear model. The lower 0.025 and upper 0.975 percentiles of those data give lower and upper values of the 95% confidence intervals.

| Noise | *p* | Threshold | Bat | Lower 95% CI | Upper 95% CI |
| --- | --- | --- | --- | --- | --- |
| Silent (Control) | 0.7 | 7.05 | A | 5.57 | 8.37 |
| Silent (Control) | 0.7 | 5.39 | B | 4.42 | 6.41 |
| Silent (Control) | 0.7 | 7.58 | C | 5.47 | 9.81 |
| Silent (Control) | 0.7 | 11.42 | D | 8.02 | 15.14 |
| Non-overlapping | 0.7 | 5.83 | A | 4.05 | 7.49 |
| Non-overlapping | 0.7 | 3.49 | B | 2.44 | 4.63 |
| Non-overlapping | 0.7 | 11.62 | C | 6.16 | 25.07 |
| Non-overlapping | 0.7 | 12.14 | D | 9.37 | 14.98 |
| Overlapping | 0.7 | 6.91 | A | 4.59 | 9.15 |
| Overlapping | 0.7 | 6.60 | B | 4.81 | 8.38 |
| Overlapping | 0.7 | 22.92 | C | 11.77 | 702.42 |
| Overlapping | 0.7 | 621.33 | D | 25.87 | >100000 |
| Sparse OL | 0.7 | 7.63 | A | 4.97 | 10.81 |
| Sparse OL | 0.7 | 4.54 | B | 2.85 | 5.95 |
| Sparse OL | 0.7 | 28.40 | C | 10.32 | >100000 |
| Sparse OL | 0.7 | 4395.17 | D | 18.24 | >100000 |

**Figure 2:**
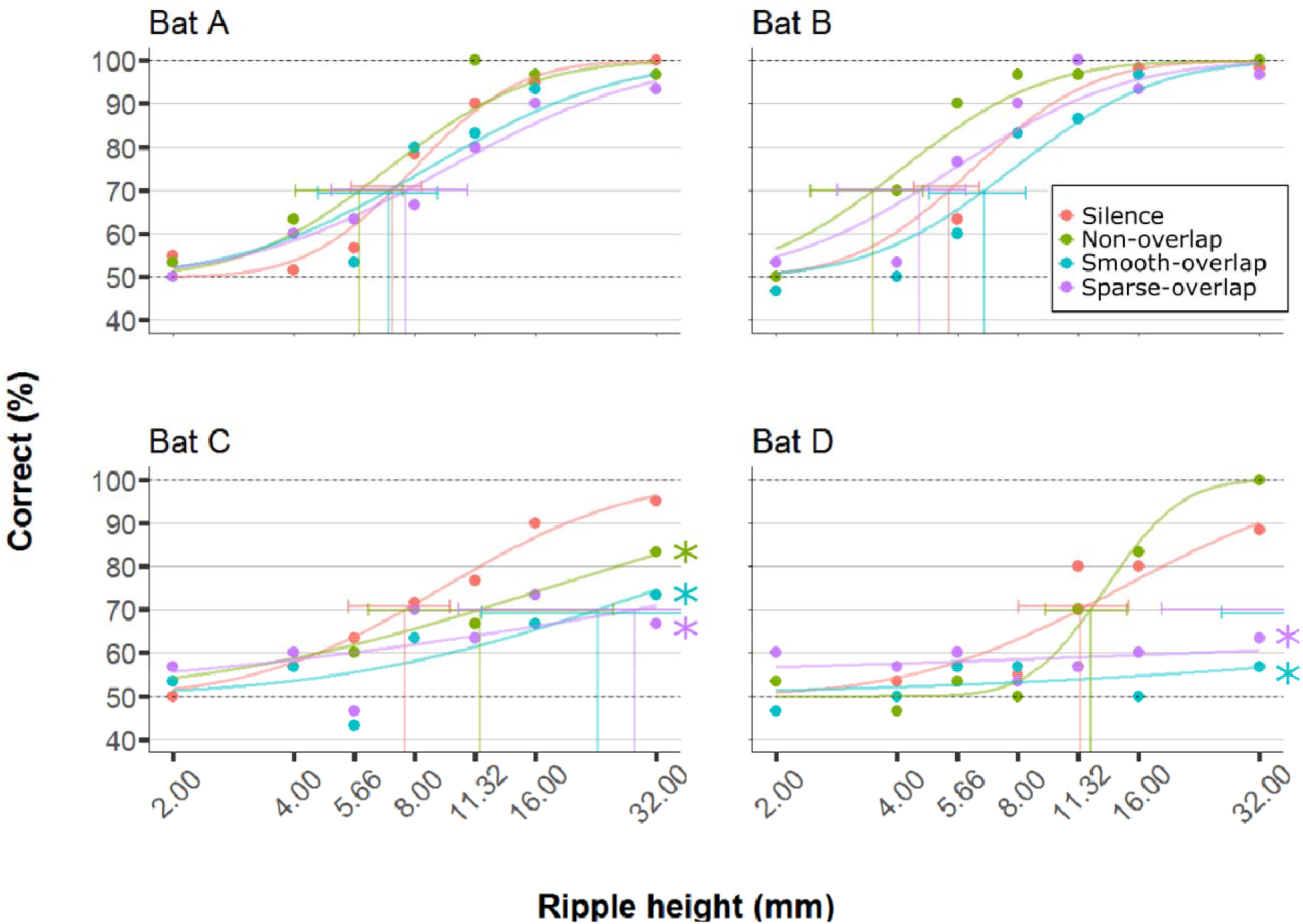
Discrimination performance of four bats as a function of peak-to-peak disc ripple height during silent control and three noise treatments. Asterisks denote that the interaction between noise treatment and ripple height (relative to silent controls; orange lines) differs significantly in generalized linear models. Discrimination performance of ‘coping’ bats (A and B) did not differ between silence and noise treatments. In contrast, discrimination performance of ‘non-coping’ bats (C and D) was reduced under both overlapping noise treatments (blue and purple lines), while discrimination performance of bat C was also reduced in non-overlapping smooth noise (green lines).

### Trial duration

The time to complete trials differed between some noise treatments for some bats (**Figure 3**). Both bats A and D made faster decisions during smooth-overlapping noise compared to silence (model estimated trial durations of bat A and D under noise and silence, respectively: 28.8 s vs. 30.0 s (A) and 10.3 s vs. 14.8 s (D); z = −3.4, *p* < 0.01 (A); z = −4, *p* < 0.001 (D); **Table S1**). However, bat C took longer to complete trials during sparse-overlapping noise (48.6 s vs. 18.5 s; z = 4.8, *p* < 0.001), while noise treatments did not affect the trial time of bat B (**Table S1**).

**Figure 3:**
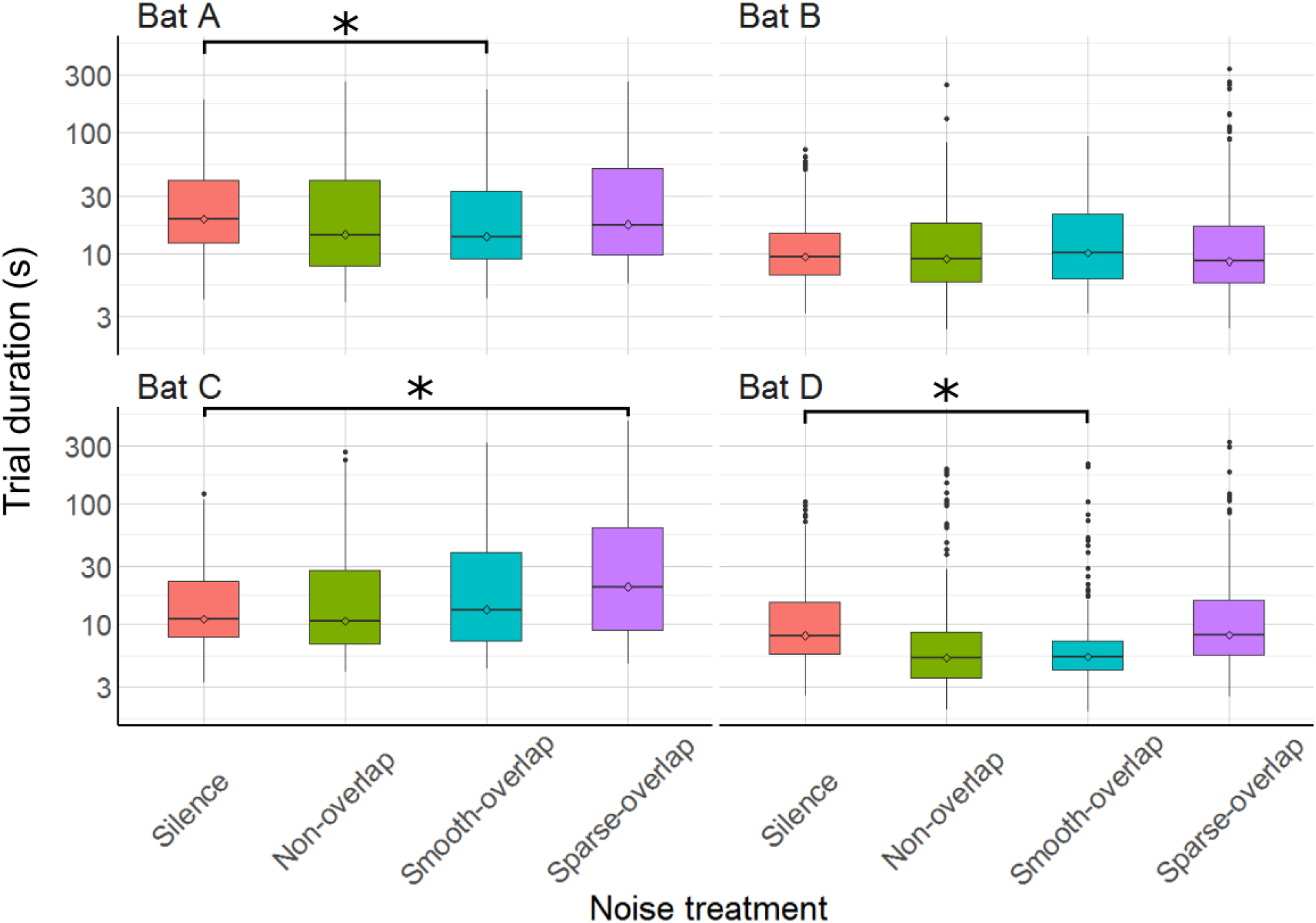
Trial duration of the discrimination task by noise treatment. Asterisks denote significant differences in trial duration relative to the control treatment. Smooth-overlapping noise reduced trial duration of bats A and D, and sparse-overlapping noise increased trial duration of bat C. Box plots show median and first and third quartiles. Whiskers represent the rest of the data minus outliers, which are shown as points.

### Aborted Trials

The bats aborted 297 trials of 4,257 total trials (7 %; bats A: 54; B: 92; C: 101; D: 50). Compared to silence, both bats B and C significantly aborted more trials under both smooth non-overlapping (z = 4.0, *p* < 0.001; z = 3.0, *p* = 0.01) and smooth-overlapping noise (z = 3.9, *p* < 0.001; z = 4.2, *p* < 0.001). In addition, bat B and also bat D aborted more trials under sparse-overlapping noise compared to silence (z = 5.2, *p* < 0.001; z = 3.8, *p* < 0.001; **Figure 4**; **Table S2**).

**Figure 4:**
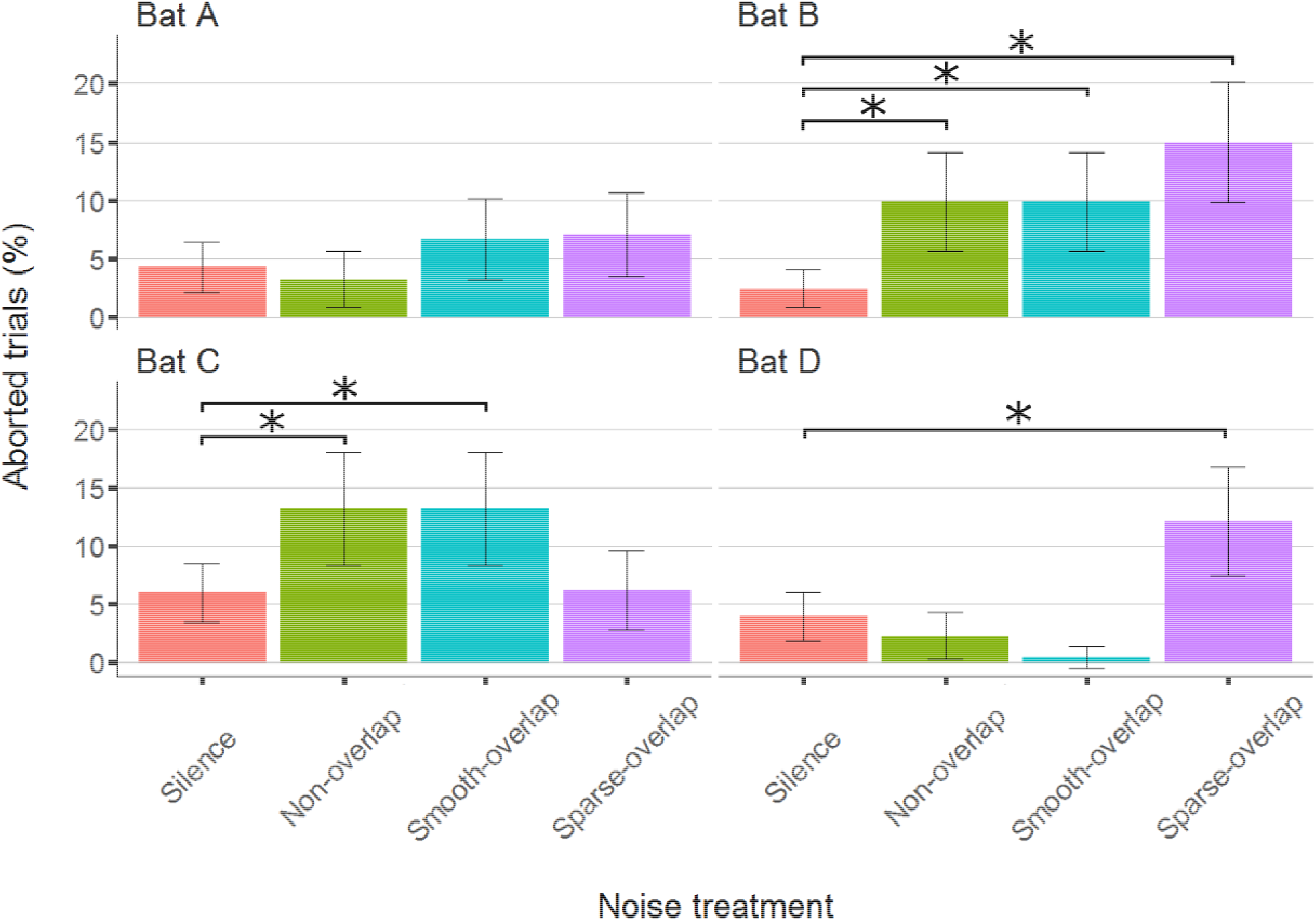
Percentage of aborted trials during the discrimination task by noise treatment. Asterisks denote significant differences in the percentage of aborted trials relative to the control treatment. Both bats B and C aborted significantly more trials for both non-overlapping and overlapping noise. Yet, bats B and D aborted significantly more trials for sparse-overlapping noise.

### Echolocation call parameters

Mean call duration ranged from 0.38 ms (bat A) to 0.47 ms (bat C). All bats increased call duration under smooth-overlapping noise. Coping bats (A and B) increased call duration by an estimated 0.07 ms, while non-coping bats (C and D) only increased call duration by 0.05 ms and 0.04 ms (bat A: t = 20.4, *p* < 0.001; B: t = 9.1, *p* < 0.001; C: t = 7.9, *p* < 0.001; D: t = 6.0, *p* < 0.001; **Table S3**). Similarly, coping bats increased call duration in sparse-overlapping noise (increase of 0.06 ms and 0.07 ms, A and B respectively), while non-coping bats did not (A: t = 15.1, *p* < 0.001; B: t = 9.3, *p* < 0.001; C: t = 1.2, *p* = 0.22; D: t = 1.8, *p* = 0.07; **Figure 5**). Oddly, bats B and C decreased call duration by 0.03 ms and 0.02 ms in non-overlapping noise relative to silence (B: t = −4.1, *p* < 0.001; C: t = −3.8, *p* < 0.001).

**Figure 5:**
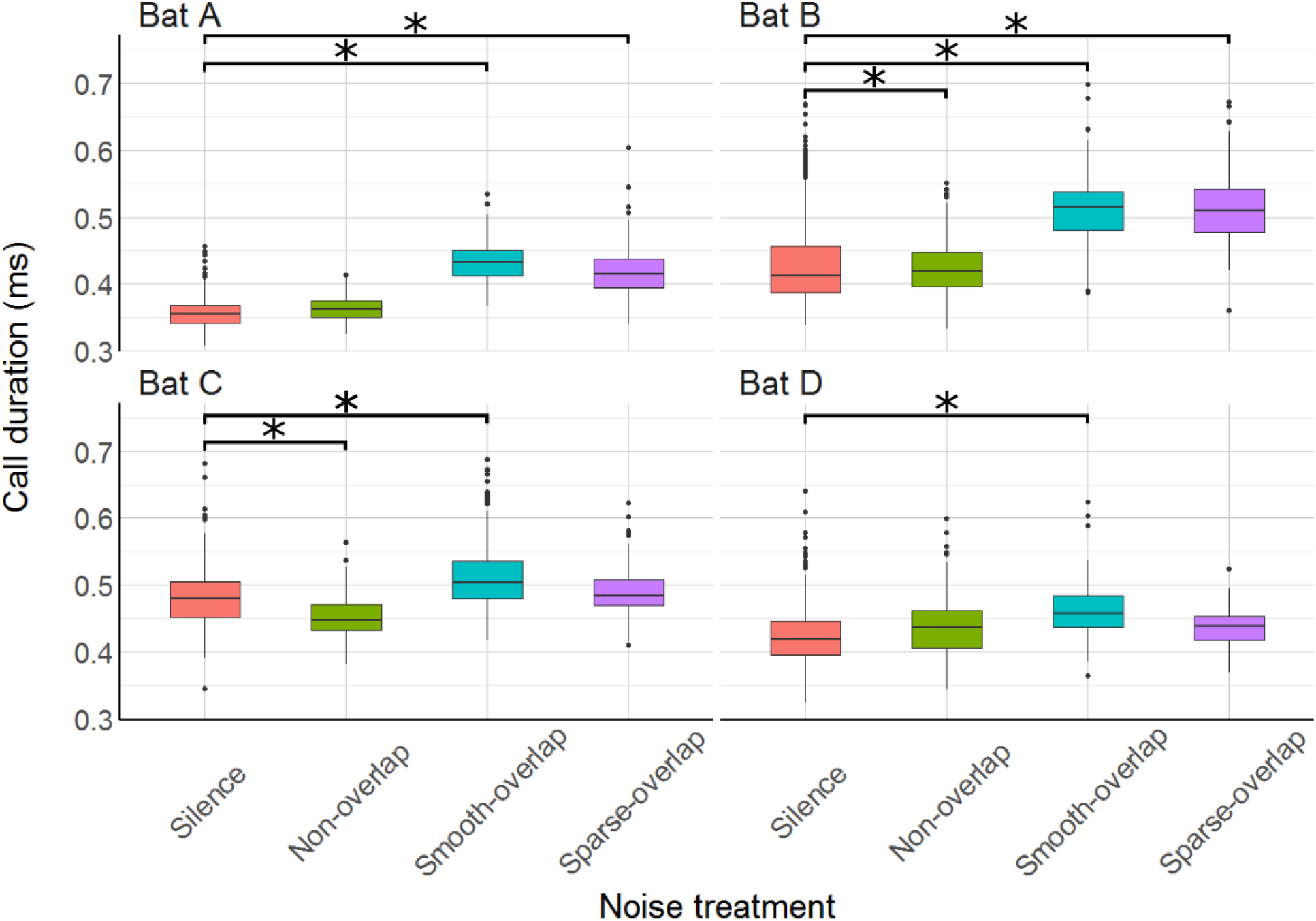
Duration of echolocation calls during the discrimination tasks by noise treatment. Asterisks indicate significant differences of call duration relative to the control treatment. All bats increased call duration under smooth-overlapping noise, and bats A and B also increased call duration under sparse-overlapping noise. Box plots show median and first and third quartiles. Whiskers represent the rest of the data minus outliers, which are shown as points.

Relative to silence, all bats increased their call sound pressure level in both overlapping noise treatments by about 10-13 dB (smooth OLN: bats A: 11.8 dB; B: 9.7 dB; C: 9.6 dB; D: 13.3 dB (t = 46.1; t = 31.8; t = 33.0; t = 25.8); sparse OLN: A: 12.6 dB; B: 8.7 dB; C: 10.5 dB; D: 13.3 dB; (t = 47.4; t = 29.8; t = 37.6; t = 19.8), all *p* < 0.001; **Figure 6**; **Table S4**). Additionally, bat A also increased call level during the smooth non-overlapping noise, though by a much lower magnitude of only 1.5 dB (t = 4.7, *p* < 0.001).

**Figure 6:**
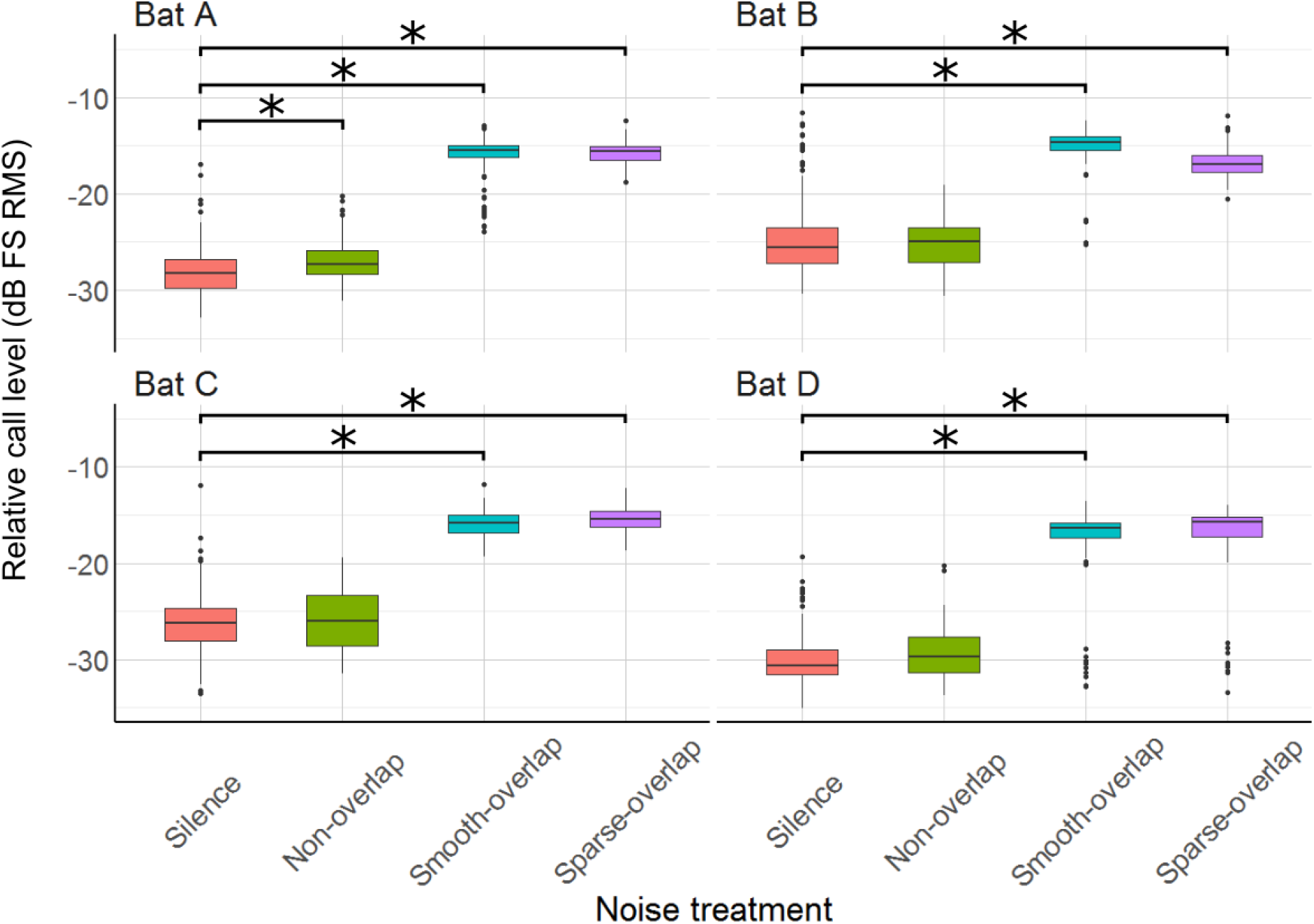
Relative sound pressure level of echolocation calls during the discrimination tasks by noise treatments. Asterisks denote significant differences relative to the control treatment. All four bats significantly increased call level under both smooth- and sparse-overlapping noise. Only bat A also increased call level under smooth non-overlapping noise, and this change was much smaller. Box plots show median and first and third quartiles. Whiskers represent the rest of the data minus outliers, which are shown as points.

The mean peak frequency was 69.8 kHz (bats A: 71.8 kHz; B: 69.9 kHz; C: 69.6 kHz; D: 67 kHz). Of all 12 comparisons, only three showed significant, yet small changes of call frequency with no clear pattern: Bat A increased peak frequency in smooth non-overlapping noise by 1.2 kHz, and decreased peak frequency in smooth overlapping noise by 1.4 kHz (t = 3.8, *p* < 0.001; t = −4.5, *p* < 0.001). Bat C increased peak frequency by 2.1 kHz only in sparse overlapping noise (t = 5.5, *p* < 0.001; **Figure 7**). Bats B and D never changed their peak frequency (**Table S5**).

**Figure 7:**
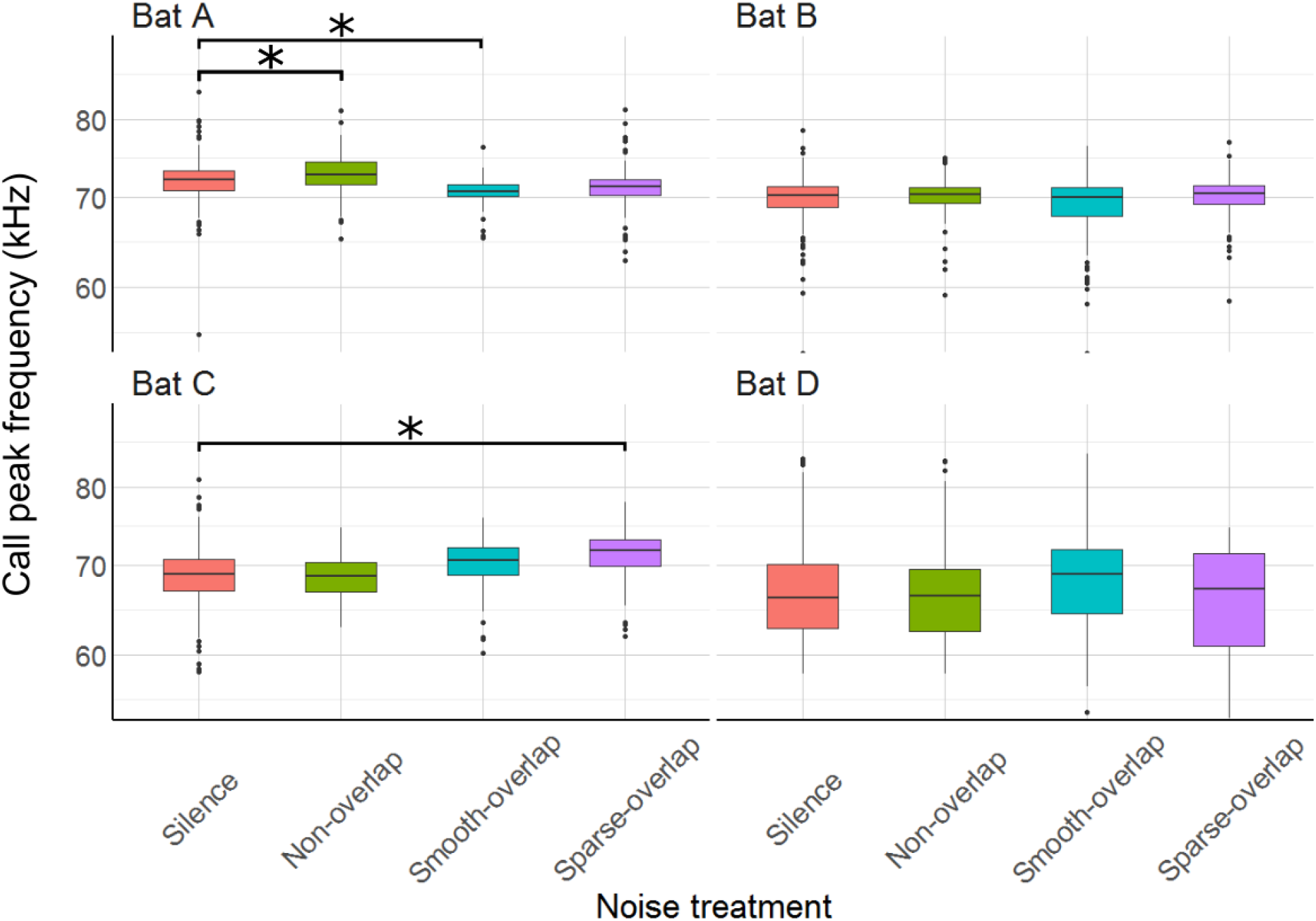
Peak frequency of echolocation calls during the discrimination tasks by noise treatment. Asterisks denote significant differences relative to the control treatment. Peak frequency of bat A increased under smooth non-overlapping noise and decreased under smooth overlapping noise. Peak frequency of bat C increased under sparse-overlapping noise. Box plots show median and first and third quartiles. Whiskers represent the rest of the data minus outliers, which are shown as points.

## Discussion

We tested the ability of four bats to discriminate increasingly rippled surface structures from a flat surface under silence and three different noise types. By comparing the bats’ discrimination performance, behavior, and echolocation parameters, we address the perceptual mechanism of noise disturbance, and how bats may be able to cope with noise disturbance. The individual bats in our experiments responded to noise in varying ways. Two bats (A and B; “coping”) were able to cope with all three noise types, as their discrimination performance was not affected by noise. In contrast, the other two bats (C and D; “non-coping”) were not able to cope with the noise, yet in different ways. Bat C had decreased discrimination performance in all three noise types, took longer in sparse-overlapping noise, and aborted more trials in smooth-overlapping and smooth non-overlapping noise. Bat D had strongly reduced discrimination performance in both smooth and sparse overlapping noise types (but not in non-overlapping noise), made faster decisions in smooth-overlapping treatments, and aborted more trials only in sparse-overlapping noise. Of the changes in echolocation call parameters, the increase in call level was the most prominent one, and shown by both coping and non-coping bats in response to both overlapping noise types. Changes in call frequency were much smaller and without a clear pattern, while call duration increased slightly more for the coping than the non-coping bats. Based on our predictions both perceptual mechanisms tested, masking and distraction, appeared to contribute to the bats’ performance. In the following, we will discuss all measured parameters in relation to our predictions about the perceptual mechanisms of noise disturbance.

### Discrimination performance (masking vs distraction)

We analyzed the ripple discrimination performance to address the perceptual mechanisms of masking and distraction. Masking should only reduce the performance in overlapping noise, and more so for smooth than sparse overlapping noise. In contrast, distraction should reduce the performance in all noise types, and most so for sparse overlapping noise. Overall, our results do not match those predictions: the coping bats (A and B) showed no decreased performance in any of the noise treatments, excluding masking and distraction. Bat C seemed to suffer from distraction, as its discrimination performance was affected by all noise types. In contrast to our prediction, however, sparse overlapping noise did not reduce performance more than the other noises. Lastly, bat D seemed to suffer from masking, as, in line with our prediction, its discrimination performance was only reduced in both overlapping noise types – yet again without difference between the smooth and sparse noise (in contrast to our prediction). The sparse noise had temporal gaps with a mean duration of 0.3 ms (range: 0-0.6 ms), which is slightly shorter than the average *P. discolor* call here (0.43 ms in silence). Although the detection performance of the gleaning bat *Megaderma lyra* for rustling sounds improved at around this gap duration (Hübner and Wiegrebe, 2003), it is possible that the temporal gaps in the sparse noise were not sufficiently long to provide sufficient release from masking for echo detection in our species *Phyllostomus discolor*. Therefore, our prediction that sparse-overlapping noise would allow bats to listen in between the gaps of the noise may be false, and further tests with larger gap widths are required. It is also possible that any release from masking that bats had gained might have been opposed by an additional distracting effect of the sparse-overlapping noise opposes, but this seems less likely than the lack of release from masking.

### Trial duration (masking vs distraction)

To further differentiate between masking and distraction as perceptual mechanisms, we also analyzed trial duration as a proxy for task difficulty. Only bat C showed a change in line with our predictions, namely a 26% increase in trial duration in sparse overlapping noise, indicative of stronger distraction by this temporally fluctuating noise. This matches our previous interpretation of this bat’s discrimination performance, suggesting that this bat was mostly affected by distraction, which should be strongest for the sparse noise. In contrast, the trial durations in smooth-overlapping noise of both the coping bat A and the non-coping bat D was even shorter than in silence, by 13 and 18%, respectively. In the coping bat A, this faster decision making did not reduce the discrimination performance, thus rather indicating reduced task difficulty due to the smooth overlapping noise, which however seems unlikely. In the non-coping bat D, the shorter trial duration might indicate frustration due to the increased task difficulty by the smooth overlapping noise. This matches our previous interpretation that this bat was affected by masking. However, it is unclear why this bat had equally reduced discrimination performance in sparse overlapping noise, but trial duration was not affected. In summary, trial duration partially supports distraction and masking as perceptual mechanisms of noise disturbance for bats C and D, respectively, but this evidence is not conclusive.

### Echolocation call characteristics (masking)

Several bat species change echolocation call parameters in response to noise (Bunkley et al., 2015; Hage et al., 2013; Luo et al., 2017; Tressler and Smotherman, 2009), which is a potential mechanism to mitigate masking effects of noise (Brumm, 2013). Thus, we next discuss whether the differences in coping ability (discrimination performance) can be explained by changes in echolocation call parameters. The most prominent change was an increase in call level by around 10-13 dB, shown by all four bats (coping and non-coping) in both overlapping noise types (smooth and sparse). This Lombard effect, the involuntary increase of vocalization amplitude in response to noise, is found in many animals from birds to humans (Brumm and Zollinger, 2011). Our species, *Phyllostomus discolor*, also exhibits an increasing Lombard effect with increasing noise level, amounting to on average +4 dB for overlapping (40-90 kHz) noise with a level of 52 dB SPL (Luo et al., 2015b). Here, we show that the Lombard effect increases even further up to 10-13 dB when noise levels are higher (70 dB SPL). This increase in call level is likely a direct response to masking (*c.f.* Fig 3 Brumm and Todt, 2002), as only one of the bats (bat A) increased call amplitude in non-masking noise, and this effect was an order of magnitude smaller (+1.5 dB, c.f. Luo et al., 2015b). Interestingly, however, although the reaction in call level was equal across all four bats, only two bats (A and B) were able to cope with masking overlapping noise in the discrimination task, while the other two bats (C and D) showed strongly reduced discrimination performance. If we assume that the increased call amplitude provides equal release from masking for all four bats, another perceptual mechanism instead of masking must be responsible for the reduced discrimination performance of the non-coping bats.

In addition to increasing call level, increased call duration improves signal detection in noise because the mammalian ear is an energy detector (Heil and Neubauer, 2003); and bats respond in this way to noise in both laboratory (Luo et al., 2015b) and field environments (Bunkley et al., 2015). Here, our bats also increased call duration, and did so only in overlapping noise types, suggesting that this was a direct response to masking. We found some differences between coping and non-coping bats. While the coping bats increased call duration by 14-16% in both overlapping noise types (smooth and sparse), the non-coping bats increased their call duration only in the smooth overlapping noise, and only by 9%. At first view, these patterns are consistent that coping bats mitigate noise masking by increasing call duration, while non-coping bats fail to do so. However, the rather small increase in call duration found here improves signal detectability by only about 1 dB (assuming a gain of 6 dB per doubling of call duration; Luo et al., 2015b). This is much less than the direct increase in call level (10-13 dB) shown by both coping and non-coping bats, making it unlikely the slight differences in call duration change can explain the differences in discrimination ability.

Shifting call frequency away from the frequency of a masker is another perceptual mechanism to improve signal detection by reducing spectral overlap, shown by bats when foraging in crowded situations (Bates et al., 2008; Gillam et al., 2007; Ratcliffe et al., 2004) or near loud ultrasonic insect choruses (Gillam and McCracken, 2007). In lower-frequency (5-35 kHz) non-overlapping noise, bat A indeed showed frequency changes consistent with avoiding spectral overlap by increasing its call peak frequency by 1.2 kHz. In contrast, the decrease of its peak frequency around 70 kHz by 1.4 kHz in the higher frequency (40-90 kHz) smooth-overlapping noise is unlikely to improve signal detectability; and correspondingly this bat did not change its peak frequency in the other overlapping noise type (sparse). Bat C increased peak frequency in sparse-overlapping noise only; and the bats B and D showed no response. It is unlikely that such small (≤ 2 kHz) changes in frequency have large effects on call detectability in noise, and thus do not seem insightful for making predictions on the ability of bats to cope with noise.

### Aborted trials

Lastly, the bats avoided the noise types differently. While the coping bat A did not abort more trials under any noise type compared to silence, the other coping bat B aborted more trials in all three noise types (6.0, 4.9, and 11.8 times more in smooth non-overlapping, smooth-overlapping, and sparse-overlapping noise, respectively). This pattern is suggestive of the noise being interpreted as danger causing fear (i.e. misleading), since the noise type did not affect the discrimination performance and trial duration in this bat (which we would expect if the bat was masked or distracted). The two non-coping bats showed opposite patterns in the number of aborted trials. Bat C aborted more trials in both smooth noise types (2.4 and 5.9 times more in smooth non-overlapping and smooth overlapping noise, respectively), but not in sparse overlapping noise. The response of bat C might indicate that smooth noise types might be interpreted as a more dangerous situation, as it cannot be linked to masking or distraction. In contrast, bat D aborted more trials in the sparse overlapping noise only (3.9 times more), but not in the two smooth noise types. It is possible here that the sparse overlapping noise was more distracting than the smooth overlapping noise, causing more trials to be aborted (somehow without affecting discrimination performance).

### Conclusion

Understanding how echolocating bats’ deal with noise pollution, as well as using noise as a deterrent to protect bats foraging near wind turbines (Arnett et al., 2013), will be important for their conservation. Recent studies have shown that echolocating bats avoid noise in the field and lab, when it is possible (Bunkley et al., 2015; Luo et al., 2015a; Schaub et al., 2008), but as noise sources expand and foraging habitat shrinks, avoidance will become more difficult. Here, when avoidance is impossible, we show that the effects of noise and the underlying perceptual mechanism of disturbance differ at the individual level. It is likely that masking affected all bats, as all of them strongly increased their call levels. However, only two of four bats were able to maintain discrimination performance in noise. Therefore, other perceptual mechanisms, in addition to masking, likely affect signal perception by bats in noise, and probably to different extents for each individual.

By grouping all individuals of one species, we may miss important differences in how individuals deal with noise (reviewed in Harding et al., 2019). By ignoring variation across individuals, we may be missing the potential for rapid evolution to occur in response to anthropogenic changes (Sih et al., 2011). Noise (or other sensory pollutants) can filter individuals over time by selecting for individuals that can cope with noise. Understanding the variation in the ability to cope with noise is paramount to predicting which species may adapt well to encroaching urbanization, and which will not. It is possible that this variation is maintained in natural systems by individual microhabitat selection, because although natural noise is ubiquitous in nature, it is spatially and temporally heterogeneous across the landscape.

## Acknowledgements

We would like to thank A. Leonie Baier for help building the experimental setup and training the bats, Karin ‘Reni’ Heckel for assistance with animal care, Yossi Yovel and Stefan Greif for help measuring light levels, and Jesse R. Barber and Henrik Brumm for comments on earlier versions of the manuscript. We thank the Max Planck Institute for Ornithology, Seewiesen, for excellent infrastructure and support, and the Fulbright Program, the National Science Foundation (GRFP ID 2018268606 to DGEG) and Deutsche Forschungsgemeinschaft (DFG, German Research Foundation, Emmy Noether grant 241711556 to HRG) for funding. All data and code are available on Zenodo (DOI: 10.5281/zenodo.3928601).

## References

Arnett, E. B., Hein, C. D., Schirmacher, M. R., Huso, M. M. and Szewczak, J. M. (2013). Evaluating the effectiveness of an ultrasonic acoustic deterrent for reducing bat fatalities at wind turbines. PloS One 8,.

Au, W. W. (2012). The sonar of dolphins. Springer Science & Business Media.

Baayen, R. H. and Milin, P. (2010). Analyzing reaction times. Int. J. Psychol. Res. 3, 12–28.

Baier, A. L., Wiegrebe, L. and Goerlitz, H. R. (2019). Echo-Imaging Exploits an Environmental High-Pass Filter to Access Spatial Information with a Non-Spatial Sensor. IScience 14, 335–344.

Barber, J. R., Crooks, K. R. and Fristrup, K. M. (2010). The costs of chronic noise exposure for terrestrial organisms. Trends Ecol. Evol. 25, 180–189.

Bates, M. E., Stamper, S. A. and Simmons, J. A. (2008). Jamming avoidance response of big brown bats in target detection. J. Exp. Biol. 211, 106–113.

Bell, G. P. and Fenton, M. B. (1986). Visual acuity, sensitivity and binocularity in a gleaning insectivorous bat, Macrotus californicus (Chiroptera: Phyllostomidae). Anim. Behav. 34, 409–414.

Boyles, J. G., Cryan, P. M., McCracken, G. F. and Kunz, T. H. (2011). Economic importance of bats in agriculture. Science 332, 41–42.

Bruintjes, R. and Radford, A. N. (2013). Context-dependent impacts of anthropogenic noise on individual and social behaviour in a cooperatively breeding fish. Anim. Behav. 85, 1343–1349.

Brumm, H. (2013). Animal communication and noise. Springer.

Brumm, H. and Slabbekoorn, H. (2005). Acoustic communication in noise. Adv. Study Behav. 35, 151–209.

Brumm, H. and Todt, D. (2002). Noise-dependent song amplitude regulation in a territorial songbird. Anim. Behav. 63, 891–897.

Brumm, H. and Zollinger, S. A. (2011). The evolution of the Lombard effect: 100 years of psychoacoustic research. Behaviour 1173–1198.

Bunkley, J. P., McClure, C. J., Kleist, N. J., Francis, C. D. and Barber, J. R. (2015). Anthropogenic noise alters bat activity levels and echolocation calls. Glob. Ecol. Conserv. 3, 62–71.

Campo, J. L., Gil, M. G. and Davila, S. G. (2005). Effects of specific noise and music stimuli on stress and fear levels of laying hens of several breeds. Appl. Anim. Behav. Sci. 91, 75–84.

Chan, A. A. Y.-H., Giraldo-Perez, P., Smith, S. and Blumstein, D. T. (2010). Anthropogenic noise affects risk assessment and attention: the distracted prey hypothesis. Biol. Lett. 6, 458–461.

Clark, C. W., Ellison, W. T., Southall, B. L., Hatch, L., Van Parijs, S. M., Frankel, A. and Ponirakis, D. (2009). Acoustic masking in marine ecosystems: intuitions, analysis, and implication. Mar. Ecol. Prog. Ser. 395, 201–222.

Dominoni, D. M., Halfwerk, W., Baird, E., Buxton, R. T., Fernández-Juricic, E., Fristrup, K. M., McKenna, M. F., Mennitt, D. J., Perkin, E. K. and Seymoure, B. M. (2020). Why conservation biology can benefit from sensory ecology. Nat. Ecol. Evol. 1–10.

Eastcott, E., Kern, J. M., Morris-Drake, A. and Radford, A. N. (2020). Intrapopulation variation in the behavioral responses of dwarf mongooses to anthropogenic noise. Behav. Ecol.

Eklöf, J., Šuba, J., Petersons, G. and Rydell, J. (2014). Visual acuity and eye size in five European bat species in relation to foraging and migration strategies. Env. Exp Biol 12, 1–6.

Esser, K.-H. and Daucher, A. (1996). Hearing in the FM-bat Phyllostomus discolor: a behavioral audiogram. J. Comp. Physiol. A 178, 779–785.

Fay, R. R. and Wilber, L. A. (1989). Hearing in vertebrates: a psychophysics databook. Acoustical Society of America.

Francis, C. D., Ortega, C. P. and Cruz, A. (2011). Noise pollution filters bird communities based on vocal frequency. PLoS One 6, e27052.

Francis, C. D., Kleist, N. J., Ortega, C. P. and Cruz, A. (2012). Noise pollution alters ecological services: enhanced pollination and disrupted seed dispersal. Proc. R. Soc. B Biol. Sci. 279, 2727–2735.

Furnham, A. and Strbac, L. (2002). Music is as distracting as noise: the differential distraction of background music and noise on the cognitive test performance of introverts and extraverts. Ergonomics 45, 203–217.

Gellermann, L. W. (1933). Chance orders of alternating stimuli in visual discrimination experiments. J. Genet. Psychol. 42, 206–208.

Gillam, E. H. and McCracken, G. F. (2007). Variability in the echolocation of Tadarida brasiliensis: effects of geography and local acoustic environment. Anim. Behav. 74, 277–286.

Gillam, E. H., Ulanovsky, N. and McCracken, G. F. (2007). Rapid jamming avoidance in biosonar. Proc. R. Soc. B Biol. Sci. 274, 651–660.

Glass, D. C. and Singer, J. E. (1972). Behavioral Aftereffects of Unpredictable and Uncontrollable Aversive Events: Although subjects were able to adapt to loud noise and other stressors in laboratory experiments, they clearly demonstrated adverse aftereffects. Am. Sci. 60, 457–465.

Goerlitz, H. R., Hübner, M. and Wiegrebe, L. (2008). Comparing passive and active hearing: spectral analysis of transient sounds in bats. J. Exp. Biol. 211, 1850–1858.

Gomes, D. G., Page, R. A., Geipel, I., Taylor, R. C., Ryan, M. J. and Halfwerk, W. (2016). Bats perceptually weight prey cues across sensory systems when hunting in noise. Science 353, 1277–1280.

Grunwald, J.-E., Schörnich, S. and Wiegrebe, L. (2004). Classification of natural textures in echolocation. Proc. Natl. Acad. Sci. 101, 5670–5674.

Hage, S. R., Jiang, T., Berquist, S. W., Feng, J. and Metzner, W. (2013). Ambient noise induces independent shifts in call frequency and amplitude within the Lombard effect in echolocating bats. Proc. Natl. Acad. Sci. 110, 4063–4068.

Halfwerk, W., Holleman, L. J. and Slabbekoorn, H. (2011). Negative impact of traffic noise on avian reproductive success. J. Appl. Ecol. 48, 210–219.

Harding, H. R., Gordon, T. A., Eastcott, E., Simpson, S. D. and Radford, A. N. (2019). Causes and consequences of intraspecific variation in animal responses to anthropogenic noise. Behav. Ecol. 30, 1501–1511.

Harrison, X. A., Donaldson, L., Correa-Cano, M. E., Evans, J., Fisher, D. N., Goodwin, C. E., Robinson, B. S., Hodgson, D. J. and Inger, R. (2018). A brief introduction to mixed effects modelling and multi-model inference in ecology. PeerJ 6, e4794.

Hartmann, W. M. and Pumplin, J. (1988). Noise power fluctuations and the masking of sine signals. J. Acoust. Soc. Am. 83, 2277–2289.

Heil, P. and Neubauer, H. (2003). A unifying basis of auditory thresholds based on temporal summation. Proc. Natl. Acad. Sci. 100, 6151–6156.

Hoffmann, S., Baier, L., Borina, F., Schuller, G., Wiegrebe, L. and Firzlaff, U. (2008). Psychophysical and neurophysiological hearing thresholds in the bat Phyllostomus discolor. J. Comp. Physiol. A 194, 39–47.

Holderied, M. W., Korine, C., Fenton, M. B., Parsons, S., Robson, S. and Jones, G. (2005). Echolocation call intensity in the aerial hawking bat Eptesicus bottae (Vespertilionidae) studied using stereo videogrammetry. J. Exp. Biol. 208, 1321–1327.

Hübner, M. and Wiegrebe, L. (2003). The effect of temporal structure on rustling-sound detection in the gleaning bat, Megaderma lyra. J. Comp. Physiol. A 189, 337–346.

Kjellberg, A., Landström, U. L. F., Tesarz, M., Söderberg, L. and Akerlund, E. (1996). The effects of nonphysical noise characteristics, ongoing task and noise sensitivity on annoyance and distraction due to noise at work. J. Environ. Psychol. 16, 123–136.

Knoblauch, K. (2007). psyphy: Functions for analyzing psychophysical data in R. R Package Version 00-5 URL HttpCRAN R-Proj. Orgpackage Psyphy.

Lattenkamp, E. Z., Vernes, S. C. and Wiegrebe, L. (2018). Volitional control of social vocalisations and vocal usage learning in bats. J. Exp. Biol. 221,.

Lazure, L. and Fenton, M. B. (2011). High duty cycle echolocation and prey detection by bats. J. Exp. Biol. 214, 1131–1137.

Luo, J., Siemers, B. M. and Koselj, K. (2015a). How anthropogenic noise affects foraging. Glob. Change Biol. 21, 3278–3289.

Luo, J., Goerlitz, H. R., Brumm, H. and Wiegrebe, L. (2015b). Linking the sender to the receiver: vocal adjustments by bats to maintain signal detection in noise. Sci. Rep. 5, 18556.

Luo, J., Lingner, A., Firzlaff, U. and Wiegrebe, L. (2017). The Lombard effect emerges early in young bats: Implications for the development of audio-vocal integration. J. Exp. Biol. 220, 1032–1037.

Matthews, K. A., Scheier, M. F., Brunson, B. I. and Carducci, B. (1980). Attention, unpredictability, and reports of physical symptoms: Eliminating the benefits of predictability. J. Pers. Soc. Psychol. 38, 525.

Naguib, M., van Oers, K., Braakhuis, A., Griffioen, M., de Goede, P. and Waas, J. R. (2013). Noise annoys: effects of noise on breeding great tits depend on personality but not on noise characteristics. Anim. Behav. 85, 949–956.

Purser, J. and Radford, A. N. (2011). Acoustic noise induces attention shifts and reduces foraging performance in three-spined sticklebacks (Gasterosteus aculeatus). PLOS One 6, e17478.

R Core Team (2017). R: A language and environment for statistical computing. Vienna, Austria: R Foundation for Statistical Computing; 2016.

Rabin, L. A., McCowan, B., Hooper, S. L. and Owings, D. H. (2003). Anthropogenic noise and its effect on animal communication: an interface between comparative psychology and conservation biology. Int. J. Comp. Psychol. 16, 172–192.

Ratcliffe, J. M., Hofstede, H. M. ter, Avila-Flores, R., Fenton, M. B., McCracken, G. F., Biscardi, S., Blasko, J., Gillam, E., Orprecio, J. and Spanjer, G. (2004). Conspecifics influence call design in the Brazilian free-tailed bat, Tadarida brasiliensis. Can. J. Zool. 82, 966–971.

Schaub, A., Ostwald, J. and Siemers, B. M. (2008). Foraging bats avoid noise. J. Exp. Biol. 211, 3174–3180.

Siemers, B. M. and Schaub, A. (2011). Hunting at the highway: traffic noise reduces foraging efficiency in acoustic predators. Proc. R. Soc. Lond. B Biol. Sci. 278, 1646–1652.

Sih, A., Ferrari, M. C. and Harris, D. J. (2011). Evolution and behavioural responses to human-induced rapid environmental change. Evol. Appl. 4, 367–387.

Simpson, S. D., Radford, A. N., Nedelec, S. L., Ferrari, M. C., Chivers, D. P., McCormick, M. I. and Meekan, M. G. (2016). Anthropogenic noise increases fish mortality by predation. Nat. Commun. 7, 10544.

Standing, L., Lynn, D. and Moxness, K. (1990). Effects of noise upon introverts and extroverts. Bull. Psychon. Soc. 28, 138–140.

Tanner Jr, W. P. (1958). What is masking? J. Acoust. Soc. Am. 30, 919–921.

Tressler, J. and Smotherman, M. S. (2009). Context-dependent effects of noise on echolocation pulse characteristics in free-tailed bats. J. Comp. Physiol. A 195, 923–934.

Tyack, P. L., Zimmer, W. M., Moretti, D., Southall, B. L., Claridge, D. E., Durban, J. W., Clark, C. W., D’Amico, A., DiMarzio, N. and Jarvis, S. (2011). Beaked whales respond to simulated and actual navy sonar. PloS One 6, e17009.

Vélez, A. and Bee, M. A. (2011). Dip listening and the cocktail party problem in grey treefrogs: signal recognition in temporally fluctuating noise. Anim. Behav. 82, 1319–1327.

Voellmy, I. K., Purser, J., Flynn, D., Kennedy, P., Simpson, S. D. and Radford, A. N. (2014). Acoustic noise reduces foraging success in two sympatric fish species via different mechanisms. Anim. Behav. 89, 191–198.

Wiley, R. H. (2013). Signal detection, noise, and the evolution of communication. In Animal communication and noise, pp. 7–30. Springer.

Zhou, Y., Radford, A. N. and Magrath, R. D. (2019). Why does noise reduce response to alarm calls? Experimental assessment of masking, distraction and greater vigilance in wild birds. Funct. Ecol. 33, 1280–1289.

